# Knowledge-guided gene prioritization reveals new insights into the mechanisms of chemoresistance

**DOI:** 10.1101/090027

**Authors:** Amin Emad, Carl R. Woese, Junmei Cairns, Krishna R. Kalari, Liewei Wang, M.D., Saurabh Sinha

**Affiliations:** Institute for Genomic Biology, University of Illinois at Urbana-Champaign, Urbana, IL 61801.; Department of Molecular Pharmacology and Experimental Therapeutics, Mayo Clinic, Rochester, MN 55905.; Department of Health Sciences Research, Mayo Clinic, Rochester, MN 55905.; Department of Computer Science and Institute of Genomic Biology, University of Illinois at Urbana-Champaign, Urbana, IL 61801.

**Keywords:** Chemoresistance, chemotherapy, drug sensitivity, gene interaction network, gene prioritization, network-based algorithm

## Abstract

**Background:** Identification of genes whose basal mRNA expression predicts the sensitivity of tumor cells to cytotoxic treatments can play an important role in individualized cancer medicine. It enables detailed characterization of the mechanism of action of drugs. Furthermore, screening the expression of these genes in the tumor tissue may suggest the best course of chemotherapy or a combination of drugs to overcome drug resistance.

**Results:** We developed a computational method called ProGENI to identify genes most associated with the variation of drug response across different individuals, based on gene expression data. In contrast to existing methods, ProGENI also utilizes prior knowledge of protein-protein and genetic interactions, using random walk techniques. Analysis of two relatively new and large datasets including gene expression data on hundreds of cell lines and their cytotoxic responses to a large compendium of drugs reveals a significant improvement in prediction of drug sensitivity using genes identified by ProGENI compared to other methods. Our siRNA knockdown experiments on ProGENI-identified genes confirmed the role of many new genes in sensitivity to three chemotherapy drugs: cisplatin, docetaxel and doxorubicin. Based on such experiments and extensive literature survey, we demonstrate that about 73% our top predicted genes modulate drug response in selected cancer cell lines. In addition, global analysis of genes associated with groups of drugs uncovered pathways of cytotoxic response shared by each group.

**Conclusions:** Our results suggest that knowledge-guided prioritization of genes using ProGENI gives new insight into mechanisms of drug resistance and identifies genes that may be targeted to overcome this phenomenon.

## BACKGROUND

The goal of gene prioritization is to rank genes with respect to their relationship to a phenotype (e.g., occurrence of a disease, response to a drug, etc.), providing an experimentalist a way to prioritize genetic perturbation tests and leading to discovery of genes affecting the phenotype [1]. In the context of drug design and drug sensitivity, various gene prioritization techniques have been used to identify drug targets, reveal mechanisms of actions (MoAs) of drugs, and identify genes associated with drug response, as well as for drug repositioning [2-5].

It has been previously shown that gene expression is the most informative currently available ‘omic’ feature with respect to drug sensitivity prediction [6], and it has been also successfully used to predict drug response in large clinical studies [7]. Basal gene expression of cancer cell lines (CCLs) has been used to rank genes by their role in cytotoxic drug resistance, utilizing correlation analysis [2, 8-11] or feature selection and regression techniques [12-16] to statistically associate drug response with gene expression profiles of cell lines. At the same time, many genes with key roles escape identification based on expression profiling alone, due to the complexity of drug MoA and noisy data [2], and due to the fact that current methods overlook known functional and biochemical relationships among genes involved in the drug MoA. Indeed, several studies have shown that utilizing such prior knowledge can improve gene prioritization based on identification of differentially expressed genes in drug-treated CCLs [3, 5, 17-19]. We posited therefore that knowledge-guided techniques should also improve analysis of basal gene expression data for identifying genes involved in drug MoA and drug sensitivity.

Although many aspects of drug MoA can be uncovered through analysis of drug-perturbed gene expression in CCLs [3, 5, 17-19], analysis of basal gene expression is valuable because it sheds light on the relationship between the cell’s resting physiological state and its drug sensitivity. In addition to the direct targets of a compound, genes and proteins involved in the processes that precede and follow the binding of the compound to its targets also play a crucial role in the compound’s MoA [20], and variations in their expression levels may underlie individual variations in drug response, even if they are not found to be differentially expressed in response to drug treatment. Thus, our primary goal here was not to identify biochemical targets of a drug or genes whose expression are affected by the drug, but rather to identify genes whose basal expression predicts the drug response. Over- or under-expression of specific genes can be experimentally shown to influence drug sensitivity [21, 22], but performing these experiments for all genes is infeasible and computational methods that can suggest candidates for such tests are necessary. Shortlisting such genes can provide complementary insight into the MoAs of a drug, offer a better understanding of drug resistance mechanisms, suggest novel targets to overcome drug resistance, and identify biomarkers of drug resistance.

We describe here a novel knowledge-guided gene prioritization algorithm called Prioritization of Genes Enhanced with Network Information (ProGENI), that discovers the relationship between basal gene expression and drug response while incorporating prior knowledge in the form of an experimentally verified network of protein-protein interactions (PPI) and genetic interactions. We used the ProGENI gene prioritization technique to analyze two large and relatively new datasets, one that includes nearly 300 human lymphoblastoid cell lines (LCLs) and another that spans over 600 CCLs of different tissues-of-origin. We employed a systematic way to evaluate different methods for gene prioritization and demonstrated the advantage of the ProGENI method. In addition, we used siRNA knockdown experiments to confirm the role of the highly ranked genes in drug sensitivity for three cytotoxic treatments widely used in chemotherapy. The results of our analysis demonstrate ProGENI to be a powerful computational technique for identifying genes that play key roles in determining drug response.

## RESULTS

### A network-based method of gene prioritization from basal expression and phenotype data

In a recent study, Rees et al. [2] identified the genes most associated with drug response variation in a collection of cell lines based on Pearson’s correlation coefficient (PCC) between basal gene expression and response, one gene at a time. We call this the ‘Pearson correlation’ scheme (or PCC) for gene prioritization. As an alternative to this ‘single gene’ analysis, we used the Elastic Net algorithm [12, 15] to perform linear regression on the drug response against the expression levels of all genes, employing regularization to enforce sparsity of features and thus learn the most relevant genes. Henceforth, we call this the ‘Elastic Net’ scheme (or EN). See Supplemental Methods in Additional file 1 for details.

We then developed a new method called Prioritization of Genes Enhanced with Network Information (ProGENI) that incorporates a network of known biological relationships among genes in the gene prioritization task. The method is illustrated in Fig. 1A. It is given a gene expression matrix with genes as columns and samples as rows, and a network with genes as nodes and inter-gene relationships as edges. It first performs a ‘network-based smoothing’ [23, 24] of the expression matrix so that the transformed expression value of a gene also reflects the activity level of the gene’s network-neighborhood (see Methods). Next, it identifies a pre-set (say *m*) number of genes with the highest correlation (both positive and negative) between the transformed expression values and the given phenotype measurements on samples; these are called the ‘response-correlated genes’ (RCGs). Then, it performs a random walk with restarts (RWR) on the network, using the genes from the previous step as the restart set, to obtain an equilibrium probability distribution on all the nodes of the network. These probabilities are then normalized with respect to a global equilibrium distribution over all gene nodes that does not depend on the RCG set. Finally, the normalized score for each node is used as the ranking criterion. This approach places the strongest RCGs at or near the top of the list, but the algorithm also makes use of prior knowledge encoded in the network.

**Figure 1:**
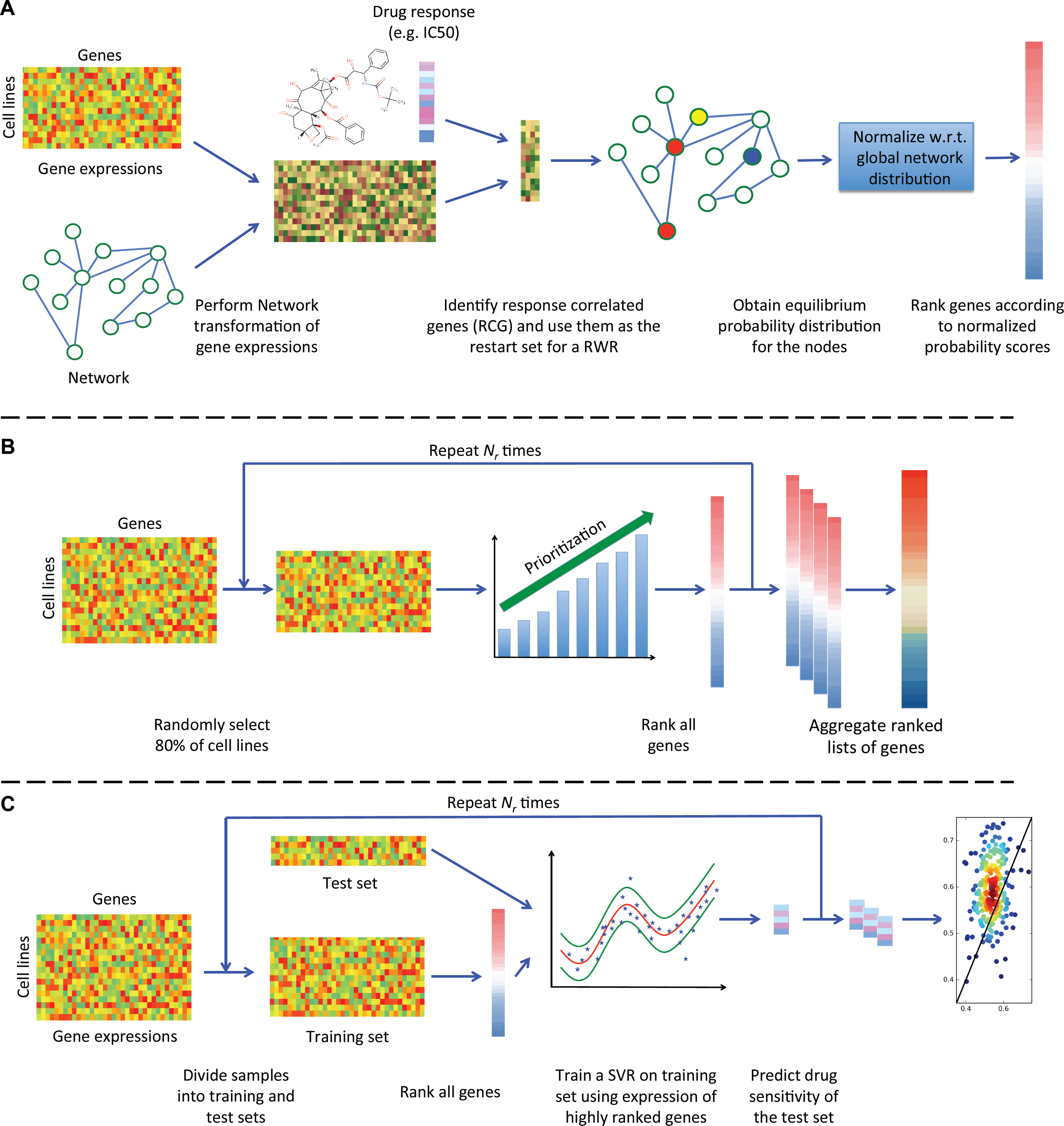
Overview of computational pipelines. (A) ProGENI: An RWR is used to obtain a vector representation of each gene and is used to perform a network transformation on the gene expressions. The response-correlated genes (RCGs) are identified as 100 genes whose transformed expressions have the highest absolute PCC with the drug response. An RWR is used to score each gene based on similarity to the RCG. These scores are then normalized to remove the network bias. (B) Robust ranking: 80% of the cell lines are selected randomly and used with a prioritization method to obtain a ranked list of genes. This procedure is repeated r times and the acquired ranked lists are aggregated to obtain a final ranked list. (C) Cross-validation scheme: a nonlinear support vector machine is trained on the training set using the top 500 genes to predict drug sensitivity of the test set and evaluate the accuracy of prediction.

To make the reported gene rankings more robust to the effect of noise in the data, we used a bootstrap sampling technique (illustrated in Fig. 1B, also see Methods), whereby prioritization is performed repeatedly on randomly selected subsets of samples and the resulting ranked lists are aggregated to produce the final ranking of genes. Henceforth, we use the name ‘Robust-ProGENI’ whenever we refer to this bootstrapping scheme and the name ‘ProGENI’ for the basic method without bootstrapping.

### Genes prioritized by ProGENI are more predictive of cytotoxic response than alternatives that do not use network information

We sought to identify the genes associated with individual variation in sensitivity to cytotoxic treatments. Towards this goal, we obtained gene expression and cytotoxic response data (EC50 values for 24 treatments) on approximately 300 LCLs from [25, 26] (see Methods). We analyzed this LCL dataset with ProGENI, using a network obtained from the STRING database [27] based on protein-protein and genetic interaction data, and focusing on one treatment at a time.

In order to evaluate the gene ranking provided by this method and other prioritization methods (Pearson correlation or Elastic Net), we used a support vector regression (SVR) algorithm to predict cytotoxic response from expression levels of the top 500 ranked genes, and assessed its accuracy with 5-fold cross-validation (see Methods and Fig. 1C). We used this evaluation setup to make sure that influential factors such as the regression algorithm and the number of used features are the same for all methods and our setup only evaluates the prioritization performance of these methods. This cross validation scheme was repeated 50 times, resulting in 250 assessments. In each assessment, the performance of the SVR was summarized using the ‘scaled probabilistic concordance index’ (SPCI) [6]. This measure ranges between 0 (bad) and 1 (good) and was specifically developed to compare drug sensitivity prediction algorithms in the DREAM 7 challenge (see Supplemental Methods in Additional file 1).

To compare the overall performance of ProGENI with baseline methods, we used the average SPCI values of the test sets for each drug (Figs. 2A and 2B). According to this evaluation scheme, drug response prediction using genes identified by ProGENI was significantly better than both the PCC-SVR scheme (FDR = 6.5E-3, one-sided Wilcoxon signed rank test on the average SPCI for each drug) and the EN-SVR scheme (FDR = 9.6E-5). We also compared the SPCI values on the 250 test sets between ProGENI and the baseline methods, separately for each treatment, using a two-sided Wilcoxon signed rank test adjusted for multiple comparisons. Since samples used in different test sets are not completely distinct, the independence assumption of the per-treatment statistical test above may be violated. Therefore, we also defined a measure called Percent of Improved Folds (PIF) as the percent of test sets for which ProGENI outperformed the baseline for any given treatment. According to these measures, ProGENI-SVR predictions were significantly better (PIF > 55%, FDR < 0.05) than the PCC-SVR for 14 (of 24) treatments and better than EN-SVR for 20 treatments; on the other hand, six treatments for PCC-SVR and three for EN-SVR showed the opposite trend (PIF < 45%, FDR < 0.05) (Table 1 and Additional file 1: Fig. S1 and Fig. S2). Fig. 3A and Fig. 3B show SPCI measures for these two prioritization schemes over all 250 test sets, for five treatments.

In addition to these evaluations, we also compared our results with two other methods: (1) EN where the number of features is not limited to top 500 and the predictions are performed using the best linear model (as opposed to using SVR) and (2) Bayesian Multitask-MKL [6], the winning method of the DREAM 7 challenge (see Supplemental Methods for details). Bayesian Multitask-MKL is a nonlinear method that uses the expression of all the genes for drug response prediction, and therefore cannot be used for gene prioritization. In spite of this, it is useful to know how well the model trained on features selected by ProGENI performs against this method. As shown in Fig. 2C and Fig. 2D, ProGENI-SVR provided significantly better predictions compared to both EN (FDR = 7.2 E-5) and Bayesian multitask-MKL (FDR = 7.2 E-5), using the average SPCI values of the test sets for each drug. In addition, ProGENI-SVR outperformed (PIF > 55%) for 20 drugs and Bayesian multitask-MKL for 22 drugs, while two drugs showed the opposite trend (PIF < 45%) for either method (Additional file 2, Additional file 1: Fig. S3 and Fig. S4). Next, we asked whether using the top 500 features identified and transformed by ProGENI improves the performance of Bayesian multitask-MKL. We found this to be the case (p-value = 8.4E-4, one-sided Wilcoxon signed rank test on the average SPCI for each drug); in addition, for 18 drugs ProGENI-Bayesian-multitask-MKL outperformed Bayesian multitask-MKL, while only for three drugs the trend was opposite (Additional file 2).

**Figure 2:**
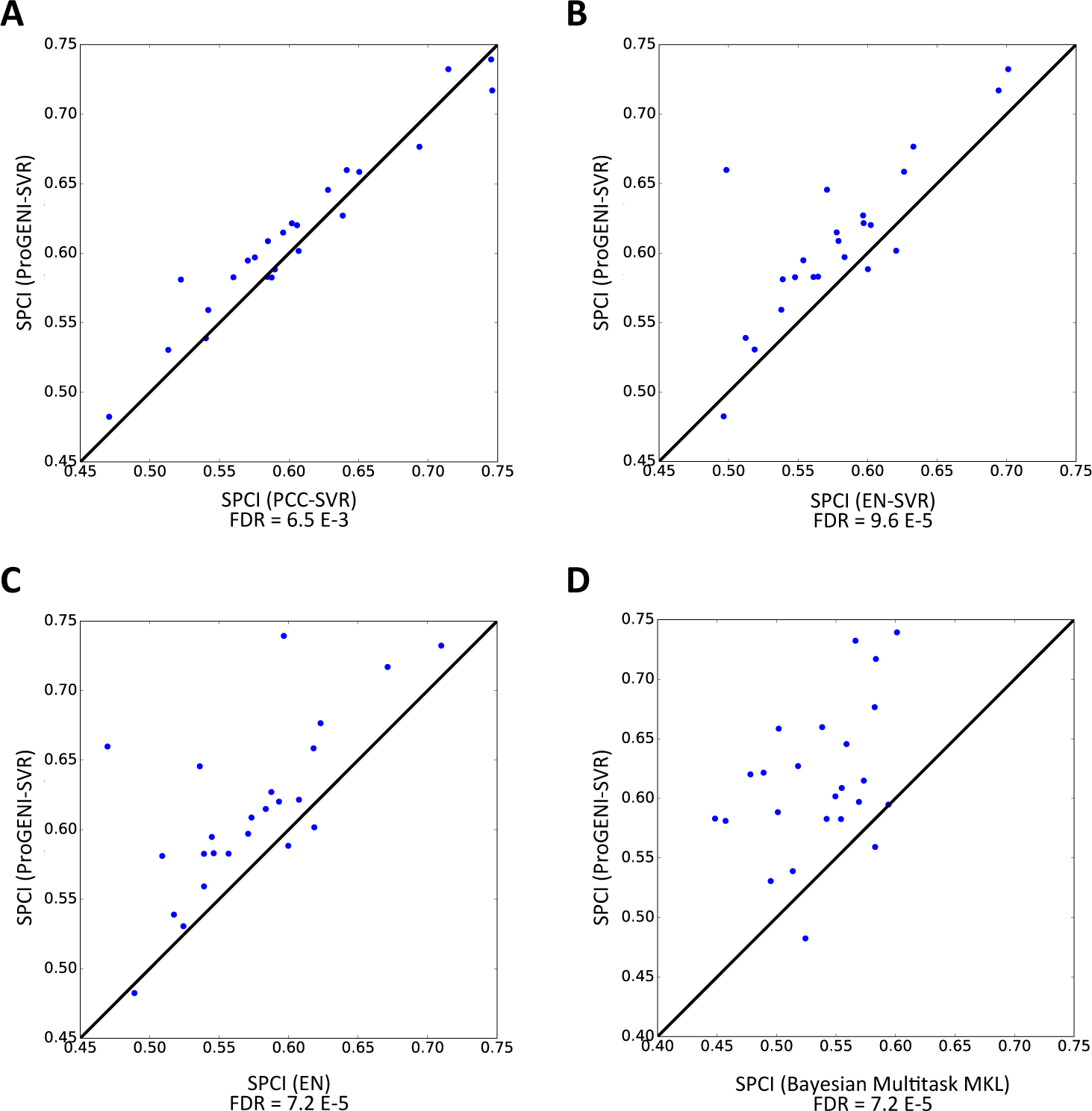
The performance of drug sensitivity prediction based on ProGENI-SVR compared to the baseline methods (PCC-SVR, EN-SVR, EN, and Bayesian multitask-MKL) using the LCL dataset for all drugs. The y-axis shows the SPCI corresponding to ProGENI-SVR while the x-axis shows the SPCI corresponding to the baseline. Each point in the scatter plot corresponds to the average SPCI value of the test sets for a single drug. The p-values are calculated using one-sided Wilcoxon signed rank test. A) Performance of ProGENI-SVR vs. PCC-SVR. B) Performance of ProGENI-SVR vs. EN-SVR. C) Performance of ProGENI-SVR vs. EN. D) Performance of ProGENI-SVR vs. Bayesian multitask-MKL.

To gain further confidence in the above observations, we proceeded to repeat the evaluation on a completely different dataset. We obtained drug response data in the form of IC50 values for 139 cytotoxic treatments and gene expression data for more than 600 CCLs from the Genomics of Drug Sensitivity in Cancer (GDSC) database from 13 tissues of origin [28]. In evaluations similar to those above, ProGENI-SVR outperformed the PCC-SVR scheme (PIF > 55%) for 66 (of 139) treatments and outperformed EN-SVR for 110 treatments (Fig. 3C and Fig. 3D, Additional file 3), while 45 and five treatments showed the opposite trend for these baseline methods, respectively. Using the average performance for each drug, evaluating performance on each drug separately, we found ProGENI-SVR to show significant improvement compared to PCC-SVR (FDR = 9.1E-4) and EN-SVR (FDR = 4.0E-21) using one-sided Wilcoxon signed rank test. ProGENI-SVR also showed significantly better performance compared to EN (FDR = 4.2E-18), with 97 drugs having PIF > 55% and only eight drugs having PIF < 45% (Additional file 3). However, the improvement of ProGENI-SVR compared to Bayesian multitask-MKL was not significant (FDR = 0.46). We note, as above, that this comparison does not compare gene prioritization performance of the two methods, since the latter method does not lend itself easily to identification of the most important (predictive) genes.

**Figure 3:**
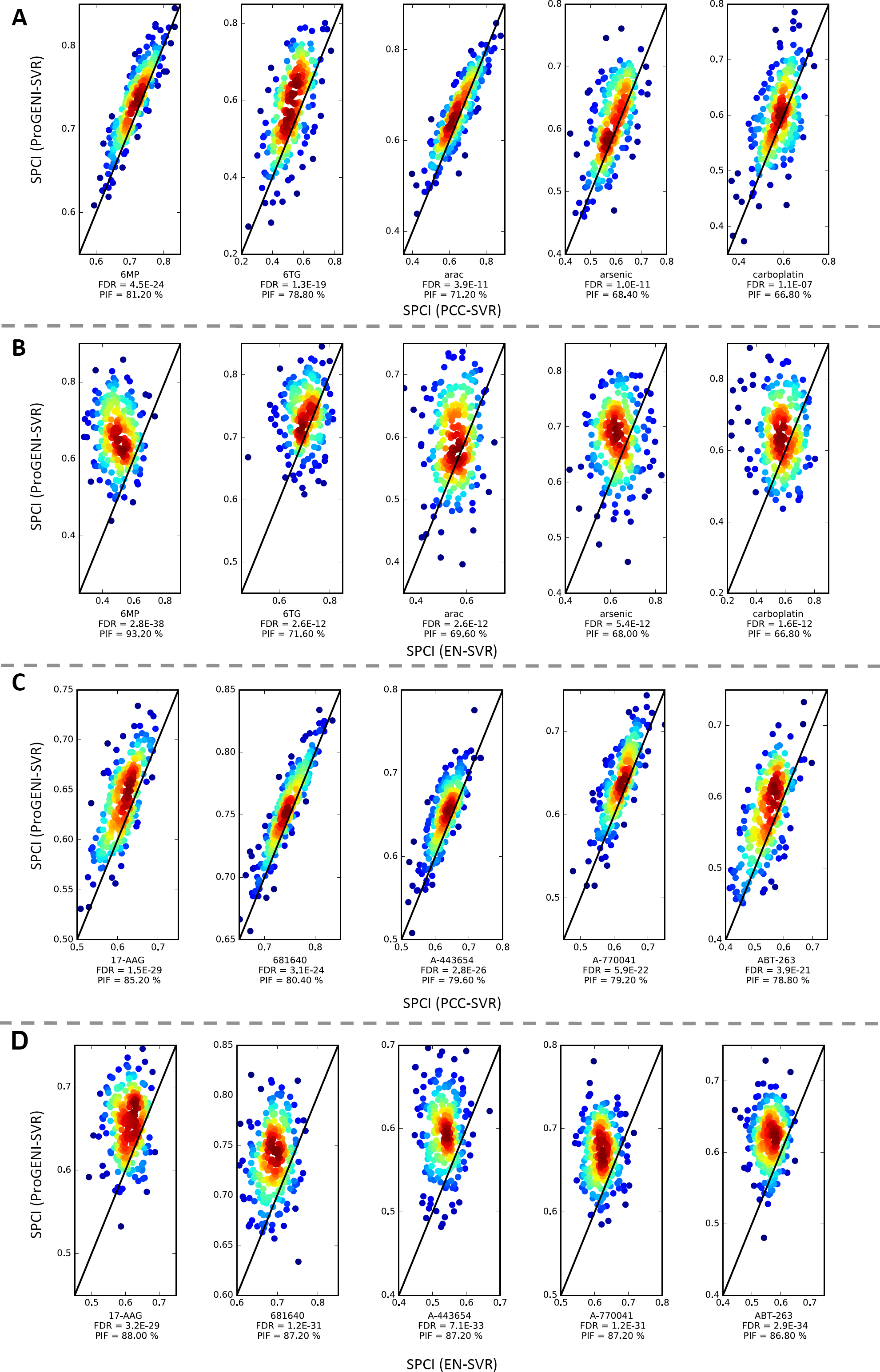
The performance of drug sensitivity prediction based on ProGENI compared to the baseline methods using the LCL (A, B) and GDSC (C, D) datasets for a selection of drugs. The y-axis shows the SPCI corresponding to ProGENI while the x-axis shows the SPCI corresponding to the baseline. Each point in the scatter plot corresponds to one random choice of training/test set. The color of each point represents the density of points in that region: a dark red color on a point means that the point is surrounded by many other points, while a blue color on a point means that the point is isolated. The FDR is calculated using a one-sided Wilcoxon signed rank test corrected for multiple tests. A) Performance of ProGENI-SVR vs. PCC-SVR for the LCL dataset. B) Performance of ProGENI-SVR vs. EN-SVR for the LCL dataset. C) Performance of ProGENI-SVR vs. PCC-SVR for the GDSC dataset. D) Performance of ProGENI-SVR vs. EN-SVR for the GDSC dataset.

**Table 1:** Performance of drug sensitivity prediction using 500 features selected by ProGENI compared to 500 features selected using baseline schemes, for the LCL dataset. The FDR is calculated using a two-sided Wilcoxon signed rank test and corrected for multiple tests. The treatments are sorted based on the largest PIF of the improvement obtained using ProGENI compared to any of the baseline schemes. Results in the range 45%<PIF<55% were considered similar. Gemcitabine was excluded, since for a few training sets the best model trained by EN only included the intercept.

| Treatment | ProGENI-SVR > PCC-SVR? (FDR, PIF) | ProGENI-SVR > EN-SVR? (FDR, PIF) | Average SPCI (ProGENI-SVR) | Average SPCI (PCC-SVR) | Average SPCI (EN-SVR) |
| --- | --- | --- | --- | --- | --- |
| doxorubicin | Yes (3.94E-11, 71.2%) | Yes (2.79E-38, 93.2%) | 0.660 | 0.641 | 0.498 |
| MPA | Yes (4.49E-24, 81.2%) | Yes (2.63E-12, 71.6%) | 0.732 | 0.714 | 0.701 |
| TCN | Yes (1.30E-19, 78.8%) | Yes (2.54E-5, 60.4%) | 0.581 | 0.522 | 0.539 |
| everolimus | Yes (1.99E-8, 66%) | Yes (2.63E-12, 69.6%) | 0.595 | 0.570 | 0.554 |
| docetaxel | Yes (1.04E-11, 68.4%) | Yes (2.84E-7, 66%) | 0.608 | 0.585 | 0.579 |
| oxaliplatin | No (1.83E-8, 32.4%) | Yes (5.44E-12, 68%) | 0.676 | 0.694 | 0.633 |
| epirubicin | Yes (2.99E-8, 65.6%) | Yes (1.61E-12, 66.8%) | 0.646 | 0.628 | 0.571 |
| NAPQI | Yes (1.11E-7, 66.8%) | Yes (1.47E-2, 56.4%) | 0.597 | 0.575 | 0.583 |
| radiation | Yes (4.30E-8, 66%) | Yes (2.38E-4, 58.8%) | 0.583 | 0.560 | 0.561 |
| carboplatin | Yes (2.66E-7, 64%) | Yes (6.67E-9, 65.2%) | 0.615 | 0.596 | 0.578 |
| MTX | No (5.52E-3, 42.4%) | Yes (2.15E-7, 64%) | 0.583 | 0.588 | 0.548 |
| cladribine | Yes (5.68E-7, 63.6%) | Yes (2.58E-5, 62.4%) | 0.622 | 0.602 | 0.597 |
| arac | Yes (5.51E-3, 60.45) | Yes (9.13E-8, 63.2%) | 0.658 | 0.650 | 0.629 |
| paclitaxel | Yes (8.94E-7, 62.8%) | Yes (6.83E-5, 61.2%) | 0.559 | 0.542 | 0.538 |
| arsenic | Yes (4.47E-6, 62.4%) | Yes (1.47E-2, 57.2%) | 0.530 | 0.513 | 0.519 |
| 6MP | No (1.59E-4, 39.2%) | Yes (2.58E-5, 62.4%) | 0.627 | 0.638 | 0.597 |
| rapamycin | Yes (3.56E-7, 61.2%) | Yes (8.23E-4, 57.6%) | 0.620 | 0.606 | 0.602 |
| 6TG | No (1.62E-19, 24.8%) | Yes (5.33E-4, 60.8%) | 0.717 | 0.746 | 0.694 |
| metformin | Similar (0.514, 48.4%) | Yes (1.87E-3, 60.4%) | 0.583 | 0.584 | 0.564 |
| fludarabine | Similar (0.655, 46.4%) | Yes (7.54E-5, 59.2%) | 0.539 | 0.540 | 0.512 |
| TMZ | Similar (2.99E-2, 54%) | No (4.23E-3, 41.6%) | 0.482 | 0.471 | 0.496 |
| hypoxia | Similar (0.406, 45.6%) | No (4.41E-2, 42.8%) | 0.588 | 0.590 | 0.600 |
| cddp | No (2.01E-3, 44%) | No (5.30E-4, 39.2%) | 0.602 | 0.607 | 0.621 |
| gemcitabine | No (1.10E-4, 37.2%) | excluded | 0.739 | 0.745 | excluded |

### Functional validations confirm the role of ProGENI-identified genes in drug response

We sought to verify whether genes associated with drug response variation (IC50 in the GDSC dataset) identified by ProGENI could be linked in vitro to significant changes in drug sensitivity. To this end, we selected the top 15 genes identified using Robust-ProGENI for three drugs – cisplatin, docetaxel, and doxorubicin – from the GDSC dataset. (These drugs belong to three different classes of cytotoxic drugs.) The selections included genes with high Pearson correlation (positive and negative) with drug response (henceforth called ‘HPC’ genes), as well as genes that were prioritized because their network neighbors’ activity was correlated with drug response. As shown in Fig. 4D, four genes for cisplatin, five genes for docetaxel, and eight genes for doxorubicin that were ranked among the top 15 by Robust-ProGENI, are not among the top 15 HPC genes. For example, the expression of *CSNK2A1*, a gene known for its role in doxorubicin response, is not highly correlated with the response to doxorubicin; however it is directly connected in the network to two HPC genes (*NOL3* and *ATF1*) and also has 23 neighbors that are directly connected to HPC genes.

**Figure 4:**
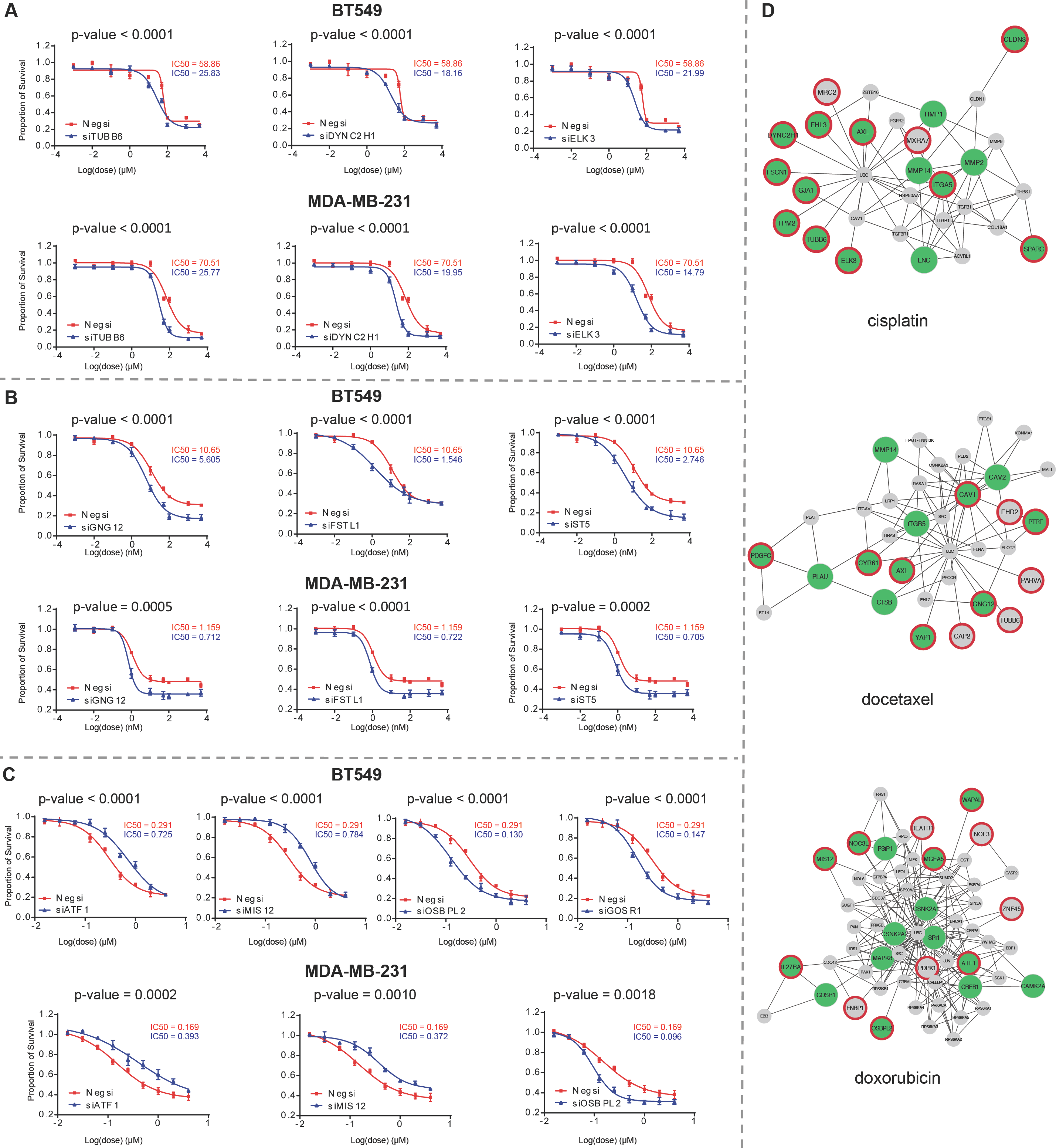
Dosage-response curves for the genes identified using Robust-ProGENI which showed significant change compared to control for (A) cisplatin, (B) docetaxel, and (C) doxorobicin in BT549 and MDA-MB-231 cell lines. Results are representative of at least three independent experiments and data expressed as mean ± standard deviation (SD), n = 3. P-values are calculated using a two-tailed unpaired t-test. D) The interaction network of genes highly ranked using Robust-ProGENI (green circles), genes highly ranked using Pearson correlation analysis (HPC) (circles with red border), and the shared neighbors of these two groups (small grey circles with no borders). Edges correspond to experimentally obtained PPI and genetic interactions extracted from STRING database; only edges with high affinity scores (>500) are depicted. The degree of a gene that is highly ranked using ProGENI but not among the HPC genes (green circle with no border) shows the number of its HPC neighbors and the number of its shared neighbors with HPC genes. These figures are drawn using Cytoscape [90].

For each identified drug-gene pair, we mined the literature for direct evidence of the gene’s role in response to that drug. Out of the 45 pairs examined, we found ‘direct’ literature evidence for 23 drug-gene pairs in that the gene’s knockdown was previously shown to affect drug sensitivity (Table 2 and Additional file 4). For predicted drug-gene pairs that were not validated by literature evidence, we performed siRNA knockdown experiments in two different cell lines of clinical significance, the human triple negative breast cancer MDA-MB-231 and BT549 cells, since these drugs are first-line therapy for triple negative breast cancer. Note that these genes were all expressed in these cell lines (Additional file 4). The siRNA knockdowns were performed for 21 candidate genes predicted by ProGENI to be associated with doxorubicin (8) docetaxel (7), or cisplatin (6), with negative siRNA as a control. The results of these assays for the 21 drug-gene pairs are shown in Fig. 4 and Additional file 1: Fig. S10 and Fig. S11, revealing that 10 of the 21 pairs were validated. Therefore, overall 33 (73%) of our 45 top predictions for these three drugs have knockdown-based evidence in their favor. Out of the top 15 genes identified using ProGENI for their role in cisplatin sensitivity, we found direct literature evidence for 9 genes (Table 2A and Additional file 4). For example, *CLDN3* (Claudin-3) (ProGENI rank 4), a gene that is involved in tight junction-specific obliteration of the intercellular space, has been shown to regulate sensitivity to cisplatin by controlling expression of cisplatin influx transporter *CTR1*; in addition, knockdown of *CLDN3* has been shown to increase resistance to cisplatin in human ovarian carcinoma cells in both in vitro culture and in vivo xenograft model [29]. As another example *MMP2*, a member of the matrix metalloproteinase family involved in the breakdown of the extracellular matrix, was ranked 9 using ProGENI, while Pearson correlation analysis did not place it among the top 15. An inhibitor of *MMP2* has been shown to significantly increase cytotoxicity in cisplatin resistant ovarian carcinoma cell line, A2780cis [30]. In addition to the 9 (of 15) genes with direct literature evidence, our own experiments revealed that knockdown of three of the remaining 6 predicted genes, *TUBB6*, *DYNC2H1*, and *ELK3*, significantly sensitized both cell lines to cisplatin treatment (Fig. 4A). β-tubulin, of which *TUBB6* is a sub-type, plays a prominent role in cell survival allowing cancer cells to survive, and these cell survival pathways can also be responsible for resistance to chemotherapy [31]. Suppression of *ELK3* induces sensitivity of MDA-MB-231 cells to doxorubicin treatment by inhibiting autophagy [32]. However, no previous study had linked these three genes to cisplatin sensitivity, making our experimental validation a novel finding.

**Table 2:**
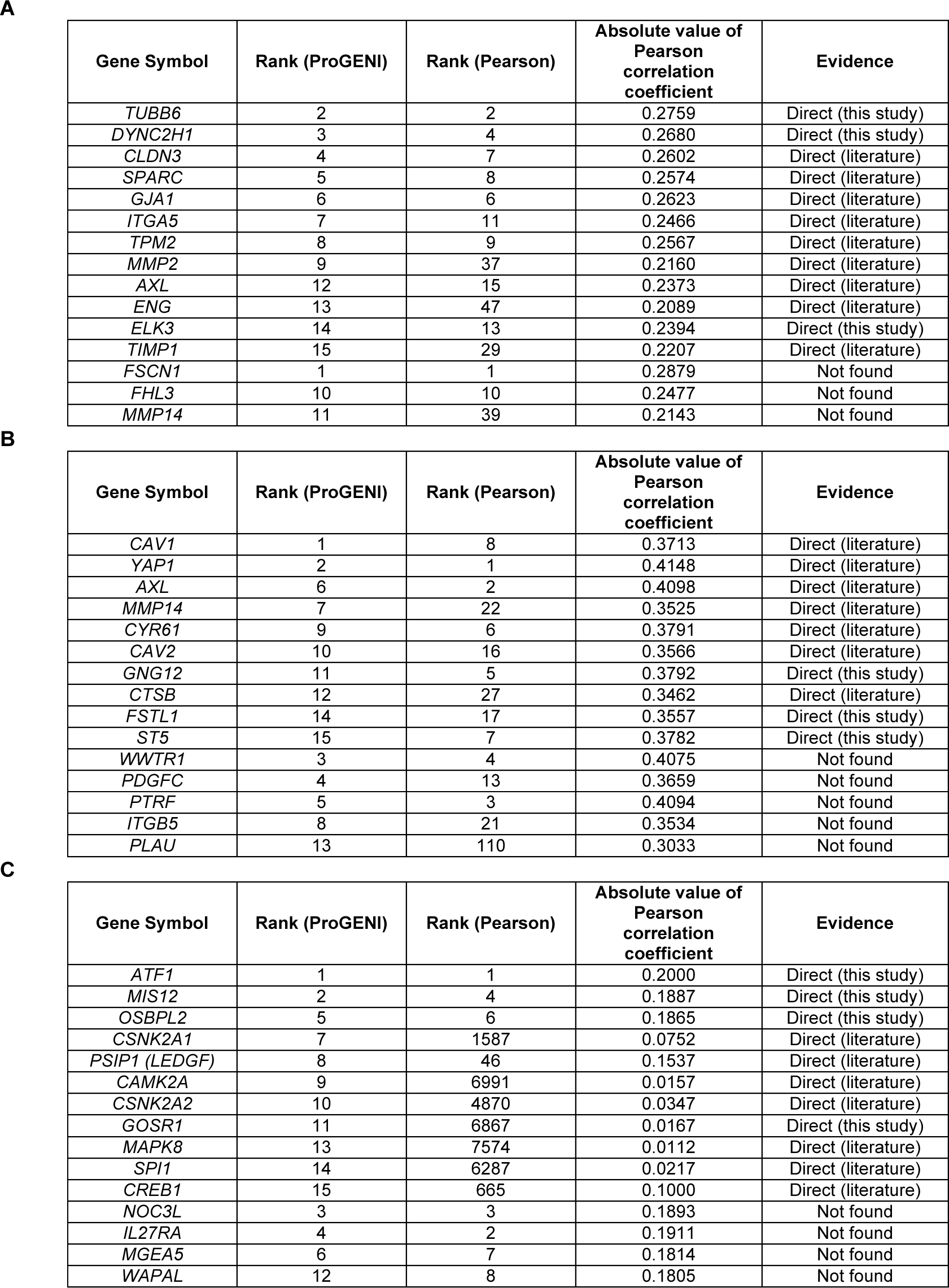
Experimental evidence for top 15 genes identified using ProGENI from the GDSC dataset for (A) cisplatin, (B) docetaxel, and (C) doxorubicin. The first column shows the gene symbols, the second column shows the rank of each gene using Robust-ProGENI, the third column shows the rank of each gene using the Pearson correlation scheme, the forth column shows the absolute value of the PCC, and the fifth column shows the nature of the evidence.

| Gene Symbol | Rank (ProGENI) | Rank (Pearson) | Absolute value of Pearson correlation coefficient | Evidence |
| --- | --- | --- | --- | --- |
| *TUBB6* | 2 | 2 | 0.2759 | Direct (this study) |
| *DYNC2H1* | 3 | 4 | 0.2680 | Direct (this study) |
| *CLDN3* | 4 | 7 | 0.2602 | Direct (literature) |
| *SPARC* | 5 | 8 | 0.2574 | Direct (literature) |
| *GJA1* | 6 | 6 | 0.2623 | Direct (literature) |
| *ITGA5* | 7 | 11 | 0.2466 | Direct (literature) |
| *TPM2* | 8 | 9 | 0.2567 | Direct (literature) |
| *MMP2* | 9 | 37 | 0.2160 | Direct (literature) |
| *AXL* | 12 | 15 | 0.2373 | Direct (literature) |
| *ENG* | 13 | 47 | 0.2089 | Direct (literature) |
| *ELK3* | 14 | 13 | 0.2394 | Direct (this study) |
| *TIMP1* | 15 | 29 | 0.2207 | Direct (literature) |
| *FSCN1* | 1 | 1 | 0.2879 | Not found |
| *FHL3* | 10 | 10 | 0.2477 | Not found |
| *MMP14* | 11 | 39 | 0.2143 | Not found |

B
| Gene Symbol | Rank (ProGENI) | Rank (Pearson) | Absolute value of Pearson correlation coefficient | Evidence |
| --- | --- | --- | --- | --- |
| *CAV1* | 1 | 8 | 0.3713 | Direct (literature) |
| *YAP1* | 2 | 1 | 0.4148 | Direct (literature) |
| *AXL* | 6 | 2 | 0.4098 | Direct (literature) |
| *MMP14* | 7 | 22 | 0.3525 | Direct (literature) |
| *CYR61* | 9 | 6 | 0.3791 | Direct (literature) |
| *CAV2* | 10 | 16 | 0.3566 | Direct (literature) |
| *GNG12* | 11 | 5 | 0.3792 | Direct (this study) |
| *CTSB* | 12 | 27 | 0.3462 | Direct (literature) |
| *FSTL1* | 14 | 17 | 0.3557 | Direct (this study) |
| *ST5* | 15 | 7 | 0.3782 | Direct (this study) |
| *WWTR1* | 3 | 4 | 0.4075 | Not found |
| *PDGFC* | 4 | 13 | 0.3659 | Not found |
| *PTRF* | 5 | 3 | 0.4094 | Not found |
| *ITGB5* | 8 | 21 | 0.3534 | Not found |
| *PLAU* | 13 | 110 | 0.3033 | Not found |

**Table 2:** Experimental evidence for top 15 genes identified using ProGENI from the GDSC dataset for (A) cisplatin, (B) docetaxel, and (C) doxorubicin.
| Gene Symbol | Rank (ProGENI) | Rank (Pearson) | Absolute value of Pearson correlation coefficient | Evidence |
| --- | --- | --- | --- | --- |
| *ATF1* | 1 | 1 | 0.2000 | Direct (this study) |
| *MIS12* | 2 | 4 | 0.1887 | Direct (this study) |
| *OSBPL2* | 5 | 6 | 0.1865 | Direct (this study) |
| *CSNK2A1* | 7 | 1587 | 0.0752 | Direct (literature) |
| *PSIP1 (LEDGF)* | 8 | 46 | 0.1537 | Direct (literature) |
| *CAMK2A* | 9 | 6991 | 0.0157 | Direct (literature) |
| *CSNK2A2* | 10 | 4870 | 0.0347 | Direct (literature) |
| *GOSR1* | 11 | 6867 | 0.0167 | Direct (this study) |
| *MAPK8* | 13 | 7574 | 0.0112 | Direct (literature) |
| *SPI1* | 14 | 6287 | 0.0217 | Direct (literature) |
| *CREB1* | 15 | 665 | 0.1000 | Direct (literature) |
| *NOC3L* | 3 | 3 | 0.1893 | Not found |
| *IL27RA* | 4 | 2 | 0.1911 | Not found |
| *MGEA5* | 6 | 7 | 0.1814 | Not found |
| *WAPAL* | 12 | 8 | 0.1805 | Not found |

Among the top 15 genes identified using ProGENI for docetaxel, we found direct literature evidence for 7 genes (Table 2B and Additional file 4). For example, *YAP1* (yes-associated protein 1) (ProGENI rank 2) regulates genes involved in cell proliferation and apoptosis; induction of this gene has been shown to induce resistance to docetaxel, and its knockdown has been shown to sensitize esophageal cancer cells to this drug [33]. Knockdowns of three of the seven remaining genes, *GNG12*, *FSTL1*, and *ST5*, significantly increased docetaxel sensitivity in both MDA-MB-231 and BT549 cells (Fig. 4B). These three genes are differentially expressed in some cancers. For example, *GNG12* is found to be down-regulated in endometrial cancer [34]. *FSTL1* was found to be downregulated in v-myc and v-ras oncogene-transformed cells, with a possible role in carcinogenesis [35], poor prognosis of glioblastoma [36], and progression of prostate cancer [37]. *ST5* (*DENND2B*) activates guanosine triphosphatase Rab13 at the leading edge of migrating cells and promotes metastatic behavior [38]. However, none of these three genes were previously known to affect docetaxel sensitivity.

We also found direct literature evidence for 8 genes among the top 15 genes for doxorubicin (Table 2C and Additional file 4). As an example, *CSNK2A1* (Casein Kinase 2 Alpha 1) and its paralog *CSNK2A2* are serine/threonine protein kinases that have regulatory roles in cell proliferation, differentiation and apoptosis. Both of these genes were ranked among the top 15 for doxorubicin using ProGENI, while Pearson correlation analysis places them at ranks 1587 and 4870, respectively. Several studies have shown the role of these genes in resistance to doxorubicin and the synergistic effect between their inhibition and cytotoxicity of doxorubicin [39-43]. As another example, Daugaard et al. have shown that the ectopic expression of *PSIP1* (*LEDGF*) (ProGENI rank 8) protects MFC-7 cells against several cytotoxic drugs including doxorubicin [44]. Through siRNA knockdown experiments, we found that three genes out of the eight remaining ‘top 15’ predictions for doxorubicin - *ATF1*, *MIS12*, and *OSBPL2* - changed doxorubicin sensitivity in both MDA-MB-231 and BT549 cells (Fig. 4C and Fig. 4D). Knockdown of *ATF1* and *MIS12* significantly desensitized both cell lines to doxorubicin treatment, while knockdown of *OSBPL2* significantly sensitized both cell lines to doxorubicin treatment. Additionally, knockdown of *GOSR1* also increased doxorubicin sensitivity in BT549 cells, but had less effect on doxorubicin response in MDA-MB-231 cells. *ATF1*, a negative regulator of apoptosis, is upregulated in metastatic melanoma cells, and inactivation of *ATF1* in melanoma cells resulted in inhibition of tumor growth and metastasis in vivo [45]. The *MIS12* complex makes an important contribution to kinetochore assembly during cell division [46]. Defects in kinetochore proteins often lead to aneuploidy and cancer. However, no previous study had linked these genes to doxorubicin sensitivity.

Finally, we note that though several of associations were not corroborated experimentally in selected breast cancer triple negative CCLs, this is expected to an extent as the selection of CCLs was based on clinical indications for each drug. In addition, the ProGENI-predicted genes were not obtained solely based on breast cancer triple negative CCLs (i.e, were not cell type specific), but rather were based on data on many different types of CCLs from 13 different tissues of origin. As a result, some of the genes that were not experimentally verified in breast cancer triple negative cell lines may influence drug response in other cell types.

### Genes highly ranked for many drugs point to common pathways of cytotoxic response

Close examination revealed that some genes are highly ranked for many treatments. Additional file 5 contains a list of 137 genes that were among the top 500 Robust-ProGENI-identified genes for at least 40 (over a quarter of 139 studied) treatments in the GDSC dataset. Functional enrichment analysis using DAVID [47]) (Additional file 5) revealed that these genes are involved in regulation of cell proliferation (43 genes, FDR = 9.19 E-18 using Fisher exact test) and regulation of cell death (28 genes, FDR = 1.67 E-5), which can be explained by the cytotoxic nature of the considered drugs. On the other hand, some of these genes encode proteins that are involved in different processes at the cell surface, such as plasma membrane (76 genes, FDR = 4.05E-08) and cell surface receptor linked signal transduction (54 genes, FDR = 2.33 E-11). Several studies have shown the involvement of plasma membrane components in multidrug-resistance (MDR) [48-50], and transport through the cell membrane, particularly vesicular transport (exosomes), has been linked to resistance to cytotoxic drugs [50]. Other enrichments include cell adhesion (45 genes, FDR = 1.11 E-21) and focal adhesion (40 genes, FDR = 5.11 E-28) and particularly the integrin family (19 genes, FDR = 3.17 E-18), which has been shown to play an important role in drug resistance [51, 52].

Seeking additional global insights about common drug-associated genes, we next formed a drugs x genes matrix indicating the top 500 genes identified for each drug, and used agglomerative clustering to identify four dense biclusters of drugs and genes that are associated with each other (Fig. 5A and Additional file 6). We performed pathway enrichment analysis on the genes in each bicluster, using DAVID (Figs. 5B-E). We noted that one of the biclusters (Fig. 5C) includes genes enriched in the MAPK signaling pathway (FDR = 6.72 E-12). This bicluster includes drugs such as ABT-263, AICAR, ATRA, bicalutamide, IPA3, lenalidomide, methotrexate, nilotinib, PAC1, Vorinostat, and VX-702 for which either the inhibition of the MAPK signaling pathway affects drug-resistance, or this pathway is involved in their MoA [53-63]. Another bicluster (Fig. 5D) includes genes enriched in the Wnt signaling pathway, and drugs such as QS11, doxorubicin, etoposide, OSU03012, thapsigargin, and tipifarnib, whose association with the Wnt signaling pathway has been confirmed in previous studies [64-69]. We also observed a bicluster (Fig. 5E) with a group of MEK inhibitors (AZD6244, CI-1040, PD-0325901, RDEA119) and genes enriched in the inflammation response pathways, consistent with prior reports of MEK inhibition resulting in anti-inflammatory response [70]. To summarize, examination of a global map of drug-gene associations predicted by ProGENI reveals subgroups of similarly acting compounds and pathways involved in their MoA.

**Figure 5:**
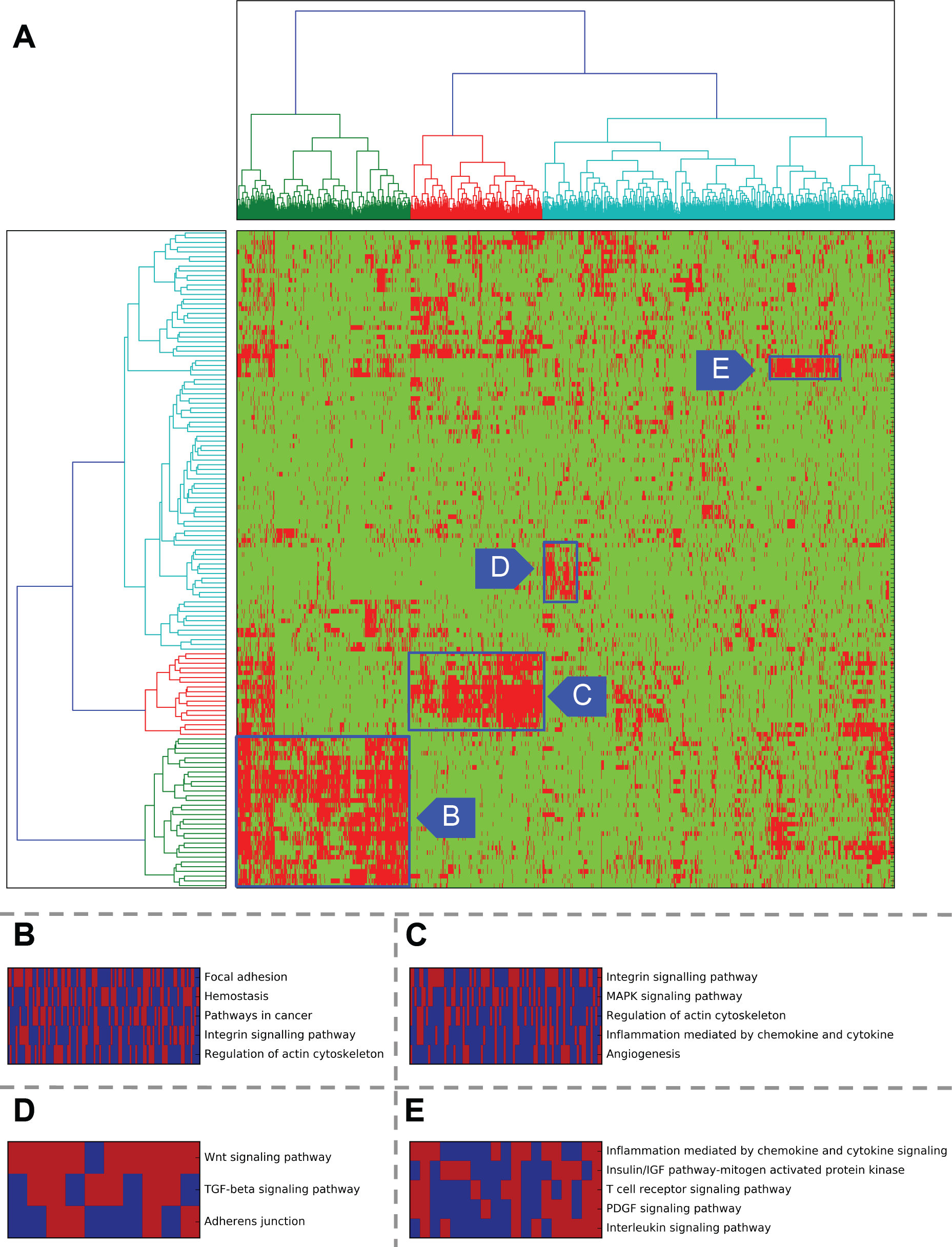
Agglomerative clustering results applied to all drugs and their Robust-ProGENI identified genes from the GDSC dataset. A) Clusters formed for drugs and genes. Rows of the matrix correspond to different drugs and columns correspond to different genes. Only columns (genes) with variance larger than 0.1 were used in the analysis (1177 genes). B, C, D, E) Enriched pathways identified for 4 cluster of genes using DAVID. Pathways with Benjamini-Hochberg corrected p-value <0.1 were sorted based on the number of shared genes and top entries were kept. Columns correspond to genes and rows correspond to pathways, with red indicating that the gene is annotated with that pathway name.

### Systematic performance analysis of ProGENI

ProGENI utilizes an interaction network consisting of different protein-protein and genetic interactions from the STRING database. To test the sensitivity of this algorithm to the choice of network, we formed a PPI network containing genetic interactions (GI), colocalizations (CO) and molecular associations (MA) from three databases: BioGRID [71], DIP [72], and IntAct [73]. The number of edges in this network (called ‘BDI’ henceforth) is approximately one third of the number of edges in the STRING network. We used the cross validation evaluation (depicted in Fig. 1C) on the LCL dataset to test the effect of changing the network to this new network. For all cases, we used SVR with Gaussian kernel as the regression algorithm to predict drug response using the identified genes. ProGENI-BDI outperformed the PCC scheme (FDR = 3.2E-2, one-sided Wilcoxon signed rank test on the average SPCI for each drug). Also, out of the 24 drugs, for 9 drugs the ProGENI-BDI performed better than PCC (PIF > 55%), while for only three drugs PCC performed better (PIF < 45%) (Fig. 6A, Additional file 1: Fig. S5, and Additional file 7). On the other hand, ProGENI-BDI had an inferior performance compared to ProGENI-STRING (FDR=4.6E-2), but their difference was small when considering number of drugs for which one method is significantly better: ProGENI-STRING outperformed ProGENI-BDI for four drugs (PIF > 55%), while ProGENI-STRING performed better for only two drugs (PIF < 45%) (Additional file 1: Fig. S6 and Additional file 7). These results show that while the choice of network plays a role in the performance of ProGENI, even a network with much smaller number of edges can significantly outperform methods that do not use the network information.

**Figure 6:**
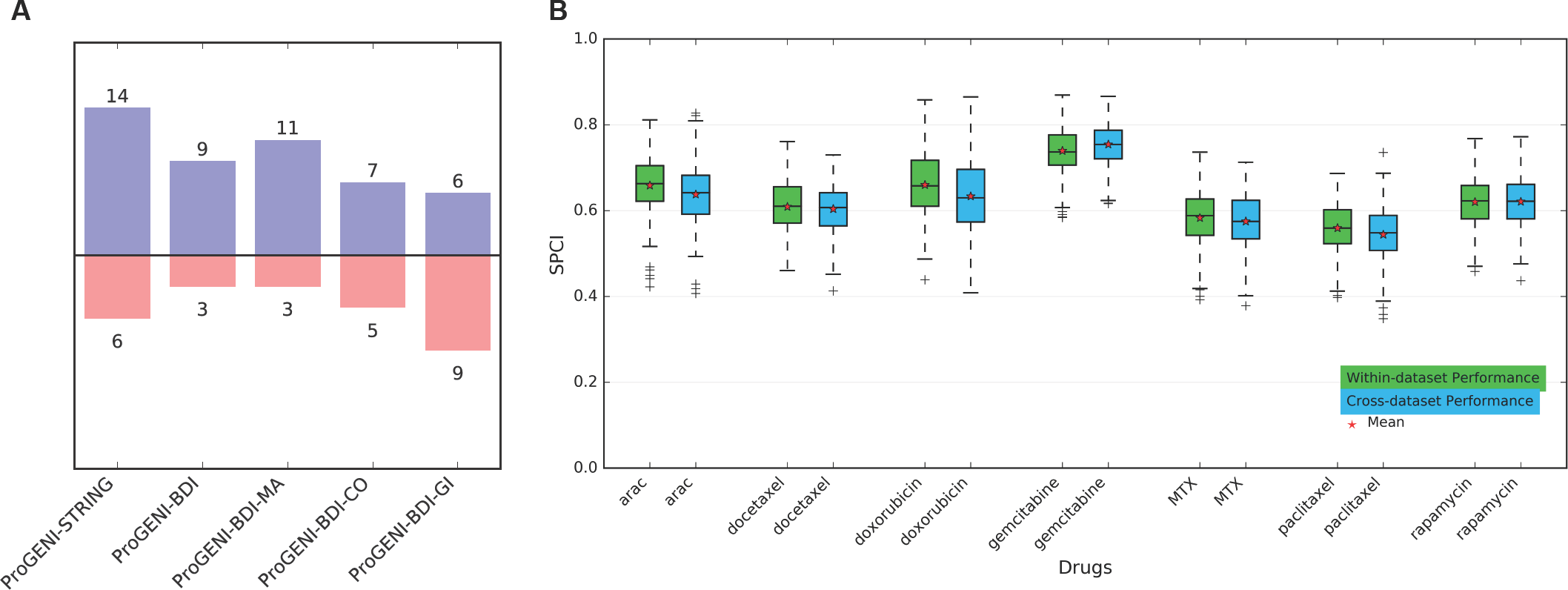
A) The performance of ProGENI-SVR with different networks compared to PCC-SVR using all 24 treatments in the LCL dataset. Color blue in each bar chart represents number of drugs for which the ProGENI-SVR performed better than the PCC-SVR (PIF > 55%), while the color red shows the number of drugs for which the trend was opposite (PIF < 45%). B) The Performance of predicting drug sensitivity of the test sets in the LCL dataset using 500 features selected by PrOGENI using the LCL dataset (within-dataset) and the GDSC dataset (cross-dataset). The box plot shows the distribution of the SPCI values for each drug.

Next, we sought to determine the role of different types of interactions on the performance of ProGENI. Of the three interaction types available in the BDI network, ProGENI with only molecular association edges had the best performance, while ProGENI with only genetic-interaction edges had the worst performance: ProGENI-BDI-MA, ProGENI-BDI-CO, and ProGENI-BDI-GI performed better than PCC on 11, 7, and 6 drugs (PIF > 55%), respectively, while PCC performed better in 3, 5, and 9 drugs (PIF < 45%), respectively (Fig. 6A, Additional file 1: Figs. S7-S9 and Additional file 7). The poor performance of ProGENI-BDI-GI may be due to the small number of edges and genes in that BDI-GI network (only ∼ 1.5 K edges among ∼1.5 K genes).

Next, we evaluated the performance of ProGENI with the STRING network to its performance when using random networks: using the LCL dataset and the STRING network, we randomly permuted the interaction network of all genes five times. In all cases, ProGENI-STRING outperformed ProGENI with the randomly permuted network (α = 0.05, one-sided Wilcoxon signed rank test on the average SPCI for each drug). ProGENI-STRING also outperformed the average performance of these five networks (p-value = 2.1E-3, one-sided Wilcoxon signed rank test on the average SPCI for each drug).

Next, we sought to study the effect of different steps in the performance of ProGENI (using the LCL dataset and the STRING network). In the first step, ProGENI performs a network transformation on the gene expression matrix, ensuring that the value assigned to each gene represents its mRNA expression and the activity level of the genes surrounding it in the network. We compared ProGENI with another variation called ‘ProGENI-PCC’ in which the absolute value of the Pearson correlation coefficient of transformed gene expressions and drug response was used to rank the genes. While the average SPCI value over all drugs was higher for ProGENI, comparing the average SPCI values for each drug did not show a significant difference (p-value = 0.42, one-sided Wilcoxon signed rank test on average SPCI values for each drug). In the second step, ProGENI selects a small set of genes (the RCG set) and scores genes in the network based on their relevance to this set. The motivation behind this step is that due to the noise in the data and the large number of genes compared to samples, it may be more reliable to score genes based on a small but high confidence set of genes. While this step showed only a slight improvement overall, for some drugs this step showed to be extremely important. For example, ProGENI significantly outperformed ProGENI-PCC for MPA (FDR = 1.95 E-19, PIF = 78%) and TCN (FDR = 8.3 E-18, PIF = 75.2%) (see Additional file 7). In summary, we found that the use of network RWR in the first and second steps improves performance, if not always with statistical significance.

We also evaluated how sensitive the performance of ProGENI is with respect to the size of the RCG set. We compared ProGENI (with an RCG set of size 100) to another variation (‘ProGENI-ACG’) in which all genes were used as the RCG set, with their restart probabilities proportional to their absolute Pearson correlation coefficient. While the average SPCI value of ProGENI was higher than ProGENI-ACG, the difference was not significant (p-value = 0.41, one-sided Wilcoxon signed rank test on average SPCI values for each drug), showing that ProGENI is not very sensitive to the size of the RCG set. However, for some drugs selecting a small RCG set had a significant effect: for example, ProGENI outperformed ProGENI-ACG for TCN (FDR = 1.3 E-19, PIF = 75.6%) and MPA (FDR = 2.32 E-22, PIF = 78.4%) (at least for some drugs) (see Additional file 7).

In light of the above evidence in favor of network-guided gene prioritization, we next asked if similarly high performance can be obtained by ignoring RCGs altogether, using only the network information. This may be possible for instance if network hubs are good predictors of drug response in general. We tested this variant method (‘NHDS’), which runs an RWR on the network with all nodes as restart set and thus prioritizes genes with high degree or genes in dense sub-networks, ignoring drug response data altogether. ProGENI significantly outperforms NHDS with p-value = 1.1 E-2 (one-sided Wilcoxon signed rank test on average SPCI values for each drug) showing that a combination of network information and information about genephenotype correlations is necessary to achieve the improved performance of ProGENI (see Additional file 7). Finally, we tested a variant (‘ProGENI-NH’) that omits the final step of adjusting for the global equilibrium distribution over gene nodes (Fig. 1A, Methods), thereby potentially advancing the ranks of network hubs. Cross validation evaluation showed ProGENI and ProGENI-NH to have very similar performance (p-value = 0.38, one-sided Wilcoxon signed rank test on average SPCI values for each drug) (see Additional file 7). However, we noted that omitting this step heavily biases the final ranked list towards network hubs (Table 3A), regardless of the phenotype being studied.

A summary of all these results is provided in Additional file 7.

### ProGENI prioritizes drug-specific genes

We noted above that a network-based prioritization method (ProGENI-NH) may show high accuracy in our cross-validation evaluation despite being heavily biased towards network hubs. If so, it is also possible that the prioritized genes are not specific to the drug being analyzed. To investigate this, we tested whether ProGENI provides a drug-specific ranking of genes. We randomly partitioned the LCL cell lines into two groups of approximately equal sizes and ran ProGENI for all drugs on these two sets. Additional file 8 reports the intersection of the top 500 genes identified for any pair of drugs using ProGENI (or Pearson correlation scheme), averaged over 100 repeats of this procedure. An expected sign of drug-specificity is that gene lists for the same drug (but based on different subsets of cell lines) have a greater intersection than gene lists for different drugs. Indeed, we noted that for 10 of the 24 treatments the intersection between gene lists based on different subsets of cell lines was ranked 1 or 2 compared to their intersection with gene lists for different drugs (Table 3B). Note that the p-value is calculated based on the fact that under the null hypothesis, the drug specificity rank follows a discrete uniform{1,24} distribution. Combing these p-values using the Fisher’s method resulted in a p-value of 2.25E-3 for drug specificity using ProGENI. This analysis shows that ProGENI provides a drug-specific set of genes. However, the results are not as specific as provided by the Pearson correlation scheme (Table 3B), which is expected since the latter relies exclusively on response data for each drug.

**Table 3:**
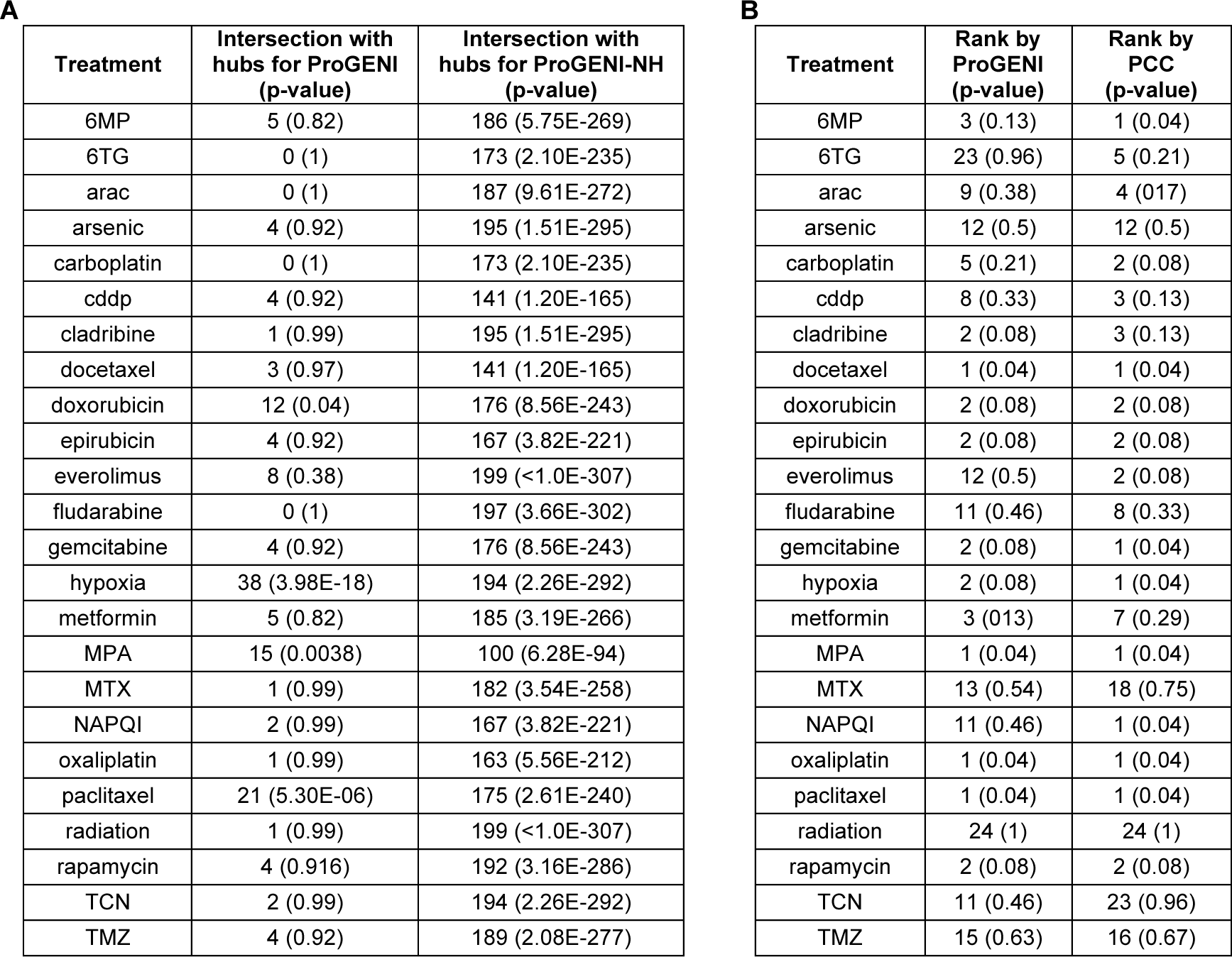
A) Presence of network hubs among the highly ranked genes provided by ProGENI and ProGENI-NH. For each treatment, this table shows the size of intersection between the top 500 genes obtained using Robust-ProGENI (or Robust-ProGENI-NH) and the set of 200 genes in the network with the highest degree. P-value for the intersection is calculated using a hypergeometric test. B) Table of drug-specificity of the top 500 genes identified using ProGENI and Pearson correlation scheme using the LCL dataset. A high rank (small entry) shows that the average size of intersection between genes identified using the prioritization method on the two sets of cell lines for the same drug is larger than the intersection when the drug is compared with other drugs. The geometric means of all ranks for prioritization using ProGENI and Pearson correlation are 4.5 and 3.1, respectively.

### Cross dataset evaluation of ProGENI

We sought to determine whether drug-associated genes identified using a heterogeneous set of cell lines (GDSC dataset) can help predict drug response in a more homogeneous cohort of cell lines (LCL dataset). We identified seven drugs shared between these two datasets, applied Robust-ProGENI on the GDSC dataset to identify top 500 genes for each drug and evaluated these genes on the LCL dataset using the SVR-based cross-validation scheme (Fig. 1C). We also compared this cross-dataset evaluation to the ‘within-dataset’ evaluation (Fig. 6B), where top genes are selected using the LCL cell lines in the training set and used to predict the drug sensitivity of the LCL cell lines in the testing set. As expected, the within-dataset evaluations yield some improvement on the performance compared to the cross-dataset evaluations, however this improvement was not statistically significant (p-value = 6.4E-2, one-sided Wilcoxon signed rank test on the average SPCI of test sets). In addition, for any given drug the average SPCI values for the two schemes were similar (Fig. 6B), suggesting that it is practical to utilize gene prioritization results from a diverse cohort such as GDSC in a more specific context such as LCLs.

## DISCUSSION

Profiling of cell lines is a promising means to better understand the mechanisms that relate the genomic and transcriptomic features to many different phenotypic outcomes [74]. In this study, we used both cancer cell lines and LCLs obtained from healthy individuals to identify genes whose over/under expression influences drug response of an individual. To achieve this goal, we proposed ProGENI, a novel method that integrates information on gene interactions and relationships with data on basal mRNA expression and drug cytotoxicity in a panel of cell lines to prioritize genes that determine drug sensitivity. We showed that genes prioritized by ProGENI can together predict drug response more accurately than top genes identified by a single gene method (Pearson correlation [2]) as well as a multiple regression method (Elastic Net). Although our main goal in this study was not to develop the best drug response prediction algorithm, and we used prediction performance only to compare different prioritization methods, we showed that ProGENI-SVR outperforms several widely used prediction algorithms. Even compared to the Bayesian multitask-MKL algorithm (the winner of the DREAM 7 challenge), our algorithm provides more accurate predictions (on the LCL dataset), or performs as well (on the GDSC dataset), with the added advantage that it can also prioritize most informative genes. We have also shown that when features selected and enhanced using ProGENI are used with the Bayesian multitask-MKL algorithm, the performance improves on the LCL dataset.

Several major steps in ProGENI differentiate it from other methods, including existing methods that utilize network information. We showed that the network transformation it performs on the gene expression matrix enables it to consider the expression of each gene in the context of its network neighbors, and greatly improves performance. The systematic removal of network bias from the output of RWR in the last step of ProGENI allows it to prioritize drug-specific genes; without this step the top ranked genes are significantly enriched in high-degree nodes (network hubs), limiting our ability to obtain a complete picture of drug resistance mechanisms, and divert our attention to generic, phenotype-independent mechanisms. Similar effects of high-degree nodes on network-based analysis have been noted before in other contexts [75]. In addition to cross-validation performance, we also used knockdown evidence from the literature and our own experiments to confirm the role of many of the ProGENI-identified genes in drug resistance. These included genes whose expression had a low correlation with drug response, but the activity of their surrounding neighbors had a high correlation.

Of the 12 genes for which siRNA knockdown did not affect drug sensitivity, eight have expression highly correlated with drug response. Since these genes have high phenotype correlation both individually and in the context of the interaction network, we speculate that the experimental validation failed because these genes are in a family of genes with similar function, and knockdown of one member is compensated by other genes in that family, or because the role of these genes in drug resistance can only be captured through their corresponding pathways, which do not become disrupted by single-gene knockdown. For example, both ProGENI and correlation analysis placed *PTRF* (Cavin-1) as an influential gene for docetaxel-resistance. Cavin-1 is a member of the cavin family proteins, which along with the caveolin family are responsible for assembly of caveolae [76]. Although our analysis showed that siRNA knockdown of *PTRF* is not sufficient to affect sensitivity to docetaxel, several studies have shown the role of caveolae and the cavin and caveolin protein families on multi drug resistance and sensitivity to various drugs including docetaxel [77-80]. We speculate therefore that the role of cavin-1 in docetaxel resistance can only be captured in the context of cavin and caveolin families and the pathways with which it is associated.

The modeling techniques we employed in this study can be extended and improved in several directions. First, in our analysis we did not consider the similarities between different treatments, and the identification of genes for each drug was performed independent of other drugs. However, we expect that many of the genes that affect drugs from the same family would be the same. As a result, incorporating drug similarity information based on their chemical structure, their known targets or known MoAs can improve prediction accuracy [81]. Another area that can potentially improve these results is incorporating genomic and epigenomic data in the analysis, as has been shown for example in [26, 82]. While gene expression data has been shown to be the most informative type of data in predicting drug response (e.g. [6]), inclusion of other types of data can provide a more comprehensive picture of genomic properties that are predictive of drug response. However, one should note that incorporating such data requires great caution, as the drastic increase in number of features necessitates a much larger number of samples to recover the signal and avoid over-fitting. While, an increase in the number of samples can be obtained by conducting comprehensive experiments and measurements on many cell lines [15], an alternative approach is to combine various datasets obtained in different studies. However, the success of this approach highly depends on consistency of the combined datasets; unfortunately, several studies have shown a lack of consistency between the drug response of large public datasets [83]. As a result, new standards and protocols may be necessary to ensure reproducibility in large-scale drug screening studies [84]. In addition, as more accurate and comprehensive datasets become available on PPI and genetic interactions among the genes, we expect that new aspects of drug resistance mechanism can be uncovered using network-based methods.

Another aspect that should be further explored in future studies is the clinical relevance of the identified genes using in vitro datasets. Our results in this study were based on in vitro cell line datasets, which have been previously shown to be able to predict the clinical drug response in vivo [7] for a few drugs. However, with the emergence of new large datasets [85, 86] which include drug response for a large cohort of drugs and samples, the role and predictive ability of genes identified in vitro can be systematically evaluated in vivo.

## CONCLUSIONS

We have shown that knowledge-guided gene prioritization using ProGENI is a powerful computational technique in identifying genes that play a key role in determining drug response and provides deeper insights into mechanisms of drug resistance that cannot be achieved otherwise. This method can be used to identify mechanistic aspects of responses to a specific drug (e.g., genes, gene sets, or pathways) and find several new candidates for experimental validation. Using this approach, for the first time we confirmed the role of 10 novel genes in sensitivity of three chemotherapy drugs. The broader applicability of this new method goes beyond pharmacogenomics studies: it offers scientists a way to identify gene predictors of any phenotype of interest while incorporating prior knowledge about genes and their mutual relationships, in a manner that the current de facto standard methods such as correlation analysis or regression analysis fail to provide.

## METHODS

### Data collection

LCL dataset: We obtained basal gene expression and drug response (half maximal effective concentration or ‘EC50’) data on 284 LCLs and 24 cytotoxic treatments from [25, 26]. GDSC dataset: We obtained gene expression and drug response (half maximal inhibitory concentration or ‘IC50’) data on 624 CCLs from 13 different tissue origins and 139 cytotoxic drugs from the Genomics of Drug Sensitivity in Cancer (GDSC) database (release-5.0) [28].

The gene interaction network used here was obtained form the STRING database [27], and consists of genetic interactions, protein associations and protein colocalizations obtained experimentally. It includes more than 1.48 M undirected weighted edges (relationships) among 15,589 nodes (genes). We also obtained a network containing genetic interactions, colocalizations and molecular associations from three databases: BioGRID [71], DIP [72], and IntAct [73]. This network (called DBI) contained approximately 294 K undirected unweighted edges among 20,788 genes. Of these edges there were ∼ 1.5 K genetic interactions among 1,478 genes, ∼ 261 K molecular associations among 20,615 genes, and ∼39 K colocalizations among 6,565 genes. Note that since two genes may be connected with more than one type of edge, we used the union of these three types of edges to find the DBI network. Additional details are in Supplemental Methods (in Additional file 1).

### Incorporating network information using random walk with restart

Several steps in the proposed algorithm ProGENI use the random walk with restart (RWR) method [87] to incorporate network information in the prioritization task. RWR is a method for quantifying the similarity between any given node of a weighted network and a given set of the nodes, called the restart set. When at a node, the walker can either move to a neighboring node, or it can jump to one of the nodes in the restart set. The probability of each of these decisions is determined by the weights of the adjacent edges and the restart probability *p*. The equilibrium probability of visiting each node in the network determines the similarity between that node and the restart set.

More formally, let *A* be an *N*_n_ × *N*_n_ symmetric adjacency matrix of the network (with *N*_n_ nodes) such that *A*(*i, j*) determines the weight of the edge between nodes *i* and *j*. Also, let *B* be the corresponding probability transition matrix obtained by normalizing each column of *A* to sum up to 1. Let *v* denote the equilibrium probability of all the nodes. This vector can be obtained iteratively using *v*^(*t*+1)^ = (1 − *p*)*Bv*^(*t*)^ + *pw*, where *w* is a probability vector of length *N*_n_ determining initial probability of restart for each node. An entry in vector *w* is equal to zero if the corresponding node is not in the restart set, and is nonzero otherwise. See Supplemental Methods (Additional file 1) for the details of the convergence criterion.

### Prioritization of genes enhanced with network information (ProGENI)

ProGENI is a method for gene prioritization that incorporates prior information on gene-gene interactions with basal gene expression and drug response data obtained from a large panel of samples (Fig. 1A). As input, this algorithm accepts a weighted undirected network of gene-gene relationships, a matrix *X* of gene expression data (samples x genes), and a vector *d* of drug response values for the samples. First, a *log*_2_ transformation followed by a Z-transform ensures that the expression of each gene across all cell lines follows a distribution with mean of zero and variance of one.

Next, a network transformation is performed on the gene expression matrix *X* to generate a ‘network-smoothed’ matrix *X’* as described next. Let *N*_n_ denote the number of nodes in the network and *N*_n_ denote the number of genes shared between the gene expression dataset and the network. For each such gene, an *N*_n_ dimensional vector representation with respect to other genes in the network is obtained using a random walk with restart (RWR). This representation is equal to the vector of equilibrium probabilities, *v*_i_, when the restart set only consists of node *i*: *w*(*i*) = 1, and *w*(*j*) = 0 for *j* ≠ *i*. Using these vector representations, an *N*_s_×*N*_s_ matrix *v* is formed, where its *i*th column is obtained from *v*_i_ by removing entries corresponding to network nodes not in the expression dataset and normalizing it to sum to 1. Finally, the network-smoothed expression matrix is obtained according to *x’* = *XV*, followed by a Z-transformation on each column.

Next, we compute for each gene *i* the absolute Pearson correlation coefficient between their network-smoothed expression (a column of *X’*) and drug response (*d*), across all samples; this is denoted by *r*_i_. Then, ‘response-correlated genes’ (RCGs) are identified as the set of *m* genes with the highest values of *r*_i_. The RCG set is used as the restart set in a RWR, in which *w*(*i*) ∝ *r*_i_ if gene *i* is an RCG. The vector *w* is scaled so that it sums to 1 and is used in a RWR to generate the equilibrium probability vector *v*_*RCG*_. In addition, a global equilibrium probability vector *v*_*global*_ is obtained by performing a RWR on the network, with the same probability of restart that was used to obtain *v*_*RCG*_, and with all the nodes as the restart set (*w*(*i*) = 1/*N*_n_ for all *i*). Finally, *v*_*RCG*_ − *v*_*global*_ is used as the ranking criterion for gene prioritization. In this study, we used a probability of restart *p* = 0.5 for all RWRs, since this value provides a good balance between the local and global topology of the network.

### Robust prioritization using bootstrap sampling and Borda rank aggregation

To obtain rankings robust to noise in the data, we used the bootstrap sampling technique (Fig. 1B). A pre-specified number of samples (80% of the cell lines) are randomly sampled, and used in the prioritization method to obtain a ranked list of genes. This procedure is repeated *N*_r_ times (a user specified number) and the geometric mean of the *N*_r_ Borda scores obtained for each gene is computed and is used as the final ranking criterion [88]. See Supplemental Methods (Additional file 1) for more details.

### Cross validation scheme for prediction of drug response

We used a cross validation scheme (Fig. 1C) to evaluate the ability of different prioritization methods in identifying genes that determine and predict drug sensitivity. We used a 5-fold cross validation procedure, repeated 50 times. In each repeat, the cell lines were randomly grouped into 5 folds; 4 folds (80% of the cell lines) were used as the training set and the remaining cell lines were used as the testing set. Prioritization methods were used to analyze gene expression of cell lines within the training set and identify 500 genes. These genes were then used to train a nonlinear support vector regression (SVR) model with Gaussian kernel, using their expression values (smoothed expression for ProGENI and original expression for baseline methods) as features. (Thus, each cell line was described by a 500 dimensional feature vector.) Hyperparameters of the SVR were learnt using a 4-fold cross validation applied inside the training set. The trained model was then used with the feature vectors corresponding to the cell lines in the test set to predict their drug sensitivity. See Supplemental Methods (in Additional file 1) for the details on the set of parameters used to train the SVR. Comparisons among methods were based on the same cross-validation partitions of cell lines.

### Cell culture and treatments for knockdown experiments

Human triple negative breast cancer MDA-MB-231 and BT549 cell lines were obtained from the American Type Culture Collection (Manassas, VA). MDA-MB-231 cells were cultured in L-15 medium containing 10% FBS at 37°C without CO2. BT549 cells were cultured in RPMI 1640 containing 10% FBS at 37°C with 5% CO2.

Doxorubicin, docetaxel, and cisplatin were purchased from Sigma-Aldrich (St. Louis, MO). Drugs were dissolved in DMSO and aliquots of stock solutions were frozen at −80°C. ONTARGETplus SMARTpool siRNAs for the candidate genes and negative control siRNA were purchased from Dharmacon to prevent off-target effects caused by both the sense and antisense strands while maintaining high silencing potency. Reverse transfection was performed for MDA-MB231and BT549 cells in 96-well plates. Specifically, 3000–4000 cells were mixed with 0.3 µL of lipofectamine RNAi-MAX reagent (Invitrogen) and 10 nM siRNA for each experiment.

Total RNA was isolated from cultured cells transfected with control or specific siRNAs with the Qiagen RNeasy kit (QIAGEN, Inc.), followed by qRT-PCR performed with the one-step, Brilliant SYBR Green qRT-PCR master mix kit (Stratagene). Specifically, primers purchased from QIAGEN were used to perform qRT-PCR using the Stratagene Mx3005P Real-Time PCR detection system (Stratagene). All experiments were performed in triplicate with beta-actin as an internal control. Reverse transcribed Universal Human reference RNA (Stratagene) was used to generate a standard curve. Control reactions lacked RNA template.

### MTS cytotoxicity assay

Cell proliferation assays were performed in triplicate at each drug concentration. Cytotoxicity assays with the lymphoblastoid were performed in triplicate at each dose. Specifically, 90 µL of cells (5 × 10^4^ cells) were plated into 96-well plates (Corning, NY) and were treated with increasing dose of specific drug or Radiation. After incubation for 72 hours, 20 µL of CellTiter 96^®^ AQueous Non-Radioactive Cell Proliferation Assay solution (Promega Corporation, Madison, WI) was added to each well. Plates were read in a Safire2 plate reader (Tecan AG, Switzerland).

Cytotoxicity assays with the tumor cell lines were performed in triplicate at each drug concentration with the CellTiter 96^®^ AQueous Non-Radioactive Cell Proliferation Assay (Promega Corporation, Madison, WI). Specifically, 90 µL of cells (5 × 10^3^ cells) were plated into 96-well plates and were treated with increasing dose of specific drug. The escalation of concentrations is provided in Supplemental Methods (Additional file 1). After incubation for 72 hours, 20 µL of CellTiter 96^®^ AQueous Non-Radioactive Cell Proliferation Assay solution (Promega Corporation, Madison, WI) was added to each well. Plates were read in a Safire2 plate reader (Tecan AG, Switzerland). Cytotoxicity was assessed by plotting cell survival versus drug concentration (on a log scale).

The p-values in Fig. 4 were calculated using a two-tailed unpaired t-test. The normality assumption for t-tests was evaluated using the Cramer-von Mises test [89] (see Additional file 1 for more details and Additional file 4 for results of this test).

## DECLARATIONS

### Availability of data and materials

We obtained gene expression and drug response corresponding to the LCL dataset from [25, 26] (accessible through GEO with accession no. GSE24277). We obtained gene expression and drug response (IC50) data on 624 CCLs from 13 different tissue origins and 139 cytotoxic drugs from the Genomics of Drug Sensitivity in Cancer (GDSC) database website (release-5.0) [28]. The STRING [27] network used in this study corresponds to “experimental” edges in the file: http://string-db.org/download/protein.links.detailed.v10/9606.protein.links.detailed.v10.txt.gz. The network from BioGRID [71] corresponds to version 3.4.143 and was downloaded from: http://thebiogrid.org/downloads/archives/Latest%20Release/BIOGRID-ALL-LATEST.mitab.zip. The network from the DIP database [72] corresponds to version dip20160731 and was downloaded from: http://dip.doe-mbi.ucla.edu/dip/script/files/2016/tab25/dip20160731.txt. Finally, the network from the IntAct database [73] was downloaded form the link below: ftp://ftp.ebi.ac.uk/pub/databases/intact/current/psimitab/intact.zip

An implementation of ProGENI and ProGENI-PCC in python, with appropriate documentation, is freely available at: https://github.com/KnowEnG/ProGENI (doi: 10.5281/zenodo.826444). ProGENI will also be available, along with the underlying network, through the cloud-based analysis framework KnowEnG (knoweng.org) upon publication.

### Ethics Approval

Not applicable.

### Authors’ contributions

AE, SS, LW conceived the study and designed the algorithm. AE implemented the algorithm and performed the statistical analyses of the algorithm. JC performed the siRNA knockdown experiments, and JC and RK performed related statistical tests. All authors contributed to the drafting of the manuscript and critical discussion of the results. All authors read and approved the final manuscript.

## Acknowledgements

This work was also supported by the Mayo Clinic Center for Individualized Medicine (CIM).

## ADDITIONAL FILES

**Additional file 1.pdf:** Includes the supplemental methods and supplemental figures.

**Additional file 2.xlsx:** Includes the performance comparison of drug sensitivity prediction based on ProGENI versus several baseline methods using the LCL dataset.

**Additional file 3.xlsx:** Includes the performance comparison of drug sensitivity prediction based on ProGENI versus several baseline methods using the GDSC dataset.

**Additional file 4.xlsx:** Includes the literature evidence for the role of top 15 genes identified for cisplatin, docetaxel and doxorubicin in drug sensitivity. Also includes mRNA expression of the genes used in experimental validations, the IC50 values of each test, and the results of test of normality in two cell lines.

**Additional file 5.xlsx:** Contains a list of 137 genes that were among the top 500 Robust-ProGENI-identified genes for at least 40 (over a quarter of 139 studied) treatments in the GDSC dataset (sheet 1). Also includes pathway and GO enrichment analysis results of this set (sheet 2).

**Additional file 6.xlsx:** List of genes and drugs in each bicluster corresponding to Fig. 4. Also includes pathway enrichment analysis results of each of these biclusters.

**Additional file 7.xlsx:** Includes the drug sensitivity prediction obtained using ProGENI with different types of network and variations of ProGENI.

**Additional file 8.xlsx:** Includes the intersection of the top 500 genes identified for any pair of drugs using ProGENI (or Pearson correlation scheme), averaged over 100 repeats of this procedure based on the LCL dataset.

